# Genome sequence and analysis of the eggplant (*Solanum melongena* L.)

**DOI:** 10.1101/824540

**Authors:** Dandan Li, Jun Qian, Wenjia Li, Yaqin Jiang, Guiyun Gan, Weiliu Li, Riyuan Chen, Ning Yu, Yan Li, Yongguan Wu, Dexian Kang, Jinmin Lian, Yongchao Niu, Yikui Wang

**Affiliations:** Institute of Vegetable Research, Guangxi Academy of Agricultural Sciences, Nanning, China; Biozeron Shenzhen, Inc., Shenzhen, China

**Keywords:** eggplant, *Solanum melongena*, genome sequencing, evolution, disease resistance, chlorogenic acid, transcription factors

## Abstract

The eggplant (*Solanum melongena* L.) is one of the most important Solanaceae crops, ranking third in the total production and economic value in the genus *Solanum*. Here, we report a high-quality, chromosome-scale eggplant reference genome sequence of 1,155.8 Mb, with N50 of 93.9 Mb, which was assembled by combining PacBio long reads and Hi-C sequencing data. Repetitive sequences occupied 70.1% of the assembly length, and 35,018 high-confidence protein-coding genes were annotated based on multiple evidence. Comparative analysis revealed 646 species-specific families and 364 positive selection genes, conferring distinguishing traits to the eggplant. We performed genome-wide identification of disease resistance genes and discovered an expanded gene family of bacterial spot resistance in the eggplant and pepper but not in tomato and potato. The genes involved in chlorogenic acid synthesis were comprehensively characterized. Highly similar chromosomal distribution patterns of polyphenol oxidase genes were observed in the eggplant, tomato, and potato genomes. The eggplant reference genome sequence will not only facilitate evolutionary studies in the Solanaceae but also facilitate their breeding and improvement.

## Introduction

Solanaceae plants are medium-sized angiosperms; they are the largest group of vegetable crops and the third largest group of economic plants. The taxa in the Solanaceae family are abundant and diverse, with 90 genera and 3,000–4,000 species. This family includes many important crop species, e.g., food crops such as potato (*Solanum tuberosum*), vegetables such as tomato (*Solanum lycopersicum*), eggplant (*Solanum melongena* L.), and pepper (*Capsicum annuum*), raw industrial materials such as tobacco (*Nicotiana tabacum*) [1, 2], and certain plant models used in research (e.g., *Nicotiana* spp., *Solanum* spp., *Petunia* spp., and *Datura* spp.) [3, 4]. Therefore, Solanaceae plants play an important role in agricultural economics and scientific research [5-8].

The eggplant, exclusively native to the Old World, belongs to the largest genus of the Solanaceae, *Solanum*, and has been listed by the Food and Agriculture Organization as the fourth largest vegetable crop. The world production of eggplants was approximately 52.3 million tons in 2017, with China being the main producer. Previous studies of the eggplant focused on the evolution [9-12], genetic linkage map [13, 14], molecular marker development [15, 16], resistance [17, 18], fruit quality [19, 20], and high-throughput genotyping [2, 21].

However, given the lack of comprehensive studies on the eggplant genome, only 775 pathogen recognition genes have been reported in the eggplant, compared to more than 1,000 genes in each of the three other Solanaceae crops (tomato, pepper, and potato) [22], which influences the progress of studies on the evolution of disease resistance in different Solanaceae plants [23]. Eggplants are the richest source of chlorogenic acid (CGA; 5-*O*-caffeoylquinic acid) [24, 25]. This dietary phenolic acid has been proven to exhibit anti-inflammatory, antimutagenic, and antiproliferative activities; however, the mechanism of CGA formation in the eggplant has not been well elucidated [26, 27]. Therefore, a high-quality reference genome is urgently needed for eggplant research. Two published eggplant references (SME_r2.5.1 and Eggplant_V3) [13, 28] were obtained by mainly employing the Illumina short-read sequencing technology, thus exhibiting assembly fragmentation and significant gap sizes.

To facilitate our understanding of the eggplant biology and evolution, we generated a chromosome-scale reference genome assembly of a cultivated eggplant variety, ‘guiqie1’, and analyzed the sequence in comparison with those of other members of the Solanaceae. Our work provides the fundamental information for unraveling the evolution and domestication of the eggplant and may ultimately lead to further improvement of this important worldwide crop.

## Results and Discussion

### Genome sequencing, assembly, and annotation

We performed genome sequencing of the eggplant with the PacBio Sequel platform using a set of 15 SMRTcells, which yielded a total of 114.5 Gb of data (average polymerase read length: 14.5 kb) (Table S1). The PacBio-only assembly contained 625 contigs, with a total length of 1,155.8 Mb and an N50 length of 5.3 Mb (maximum contig length: 21.7 Mb) (Table 1). Subsequently, we used Dovetail Hi-C data (80.7 Gb) to refine this assembly. Of the 625 contigs, 318 were sorted into 12 superscaffolds, accounting for 97.1% of the original 1,155.8-Mb assembly. The superscaffolds were further anchored to 12 linkage groups to form pseudochromosomes (Figure S1), with N50 of 93.9 Mb and a maximum length of 112 Mb (Table 1). The number of pseudochromosomes (*n* = 12) corresponded to the number of chromosomes in the eggplant and many members of the Solanaceae [29, 30].

**Table 1.**
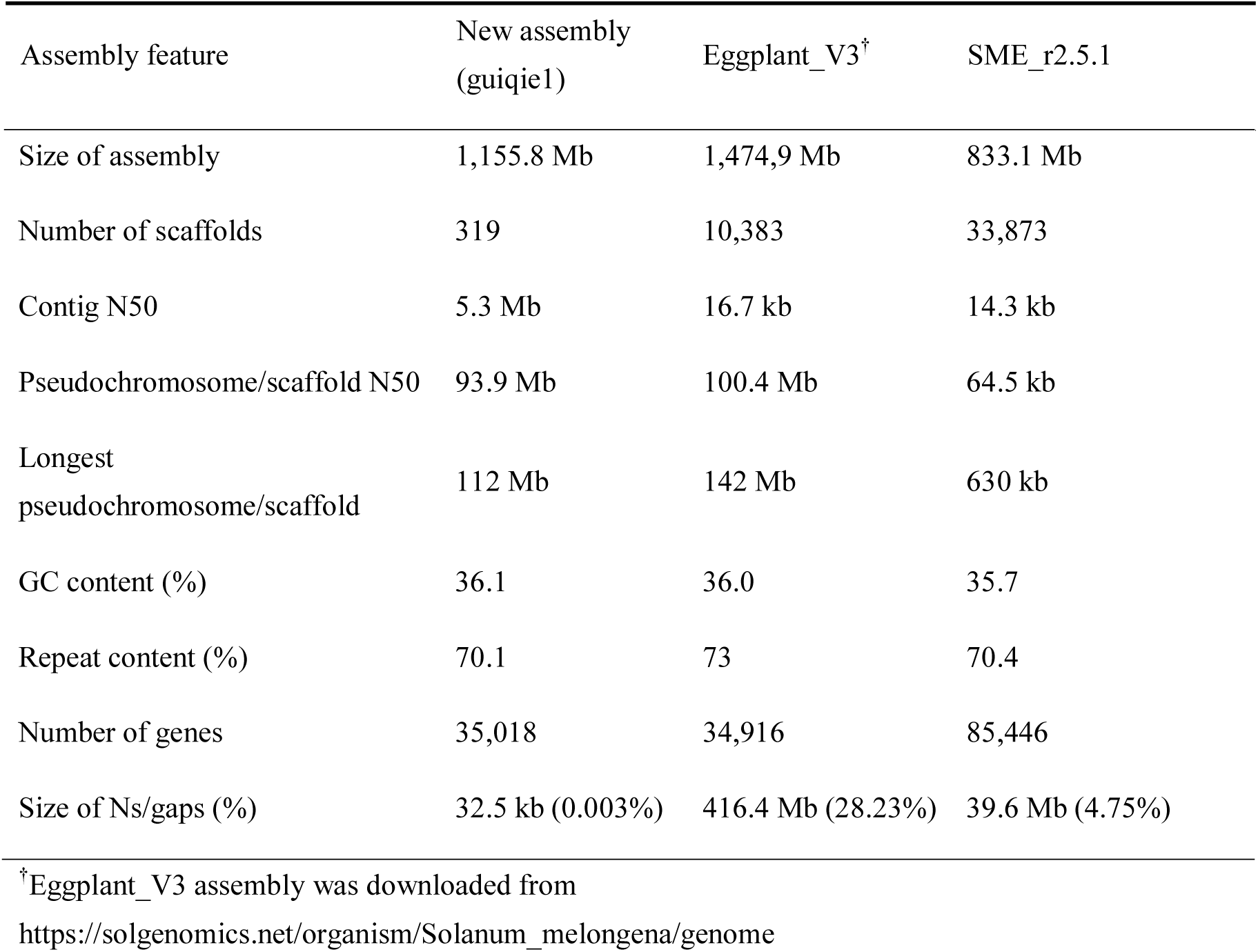
Comparison of eggplant assemblies.

Benchmarking Universal Single-Copy Ortholog (BUSCO) evaluations of the genome sequence revealed 96.2% completeness. Compared with the previously published eggplant genomes (SME_r2.5.1 and Eggplant_V3) [13, 28], which both mainly employed the Illumina short-read sequencing technology, resulting in more fragmented assemblies (contig N50 lengths: 14.3 and 16.7 kb, respectively) and larger gap sizes (Ns: 4.75% and 28.23%, respectively), our genome assembly achieved a great improvement in both quality and integrity (Table 1 and Table S2).

To validate the superscaffolds, we mapped the 952 DNA markers of linkage map LWA2010 [31] to the eggplant assembly with BWA-MEM [32] and obtained the best mapped position for each marker; a total of 946 (99.4%) markers could be mapped onto the 12 superscaffolds (Table S3). Then, ALLMAPS [33] was used with default parameters to assign the superscaffolds to each pseudochromosome, and a high value of the Pearson correlation coefficient (ρ-value > 0.9) between the physical position and map location of genetic markers indicated a high quality of the eggplant assembly (Figure S2). We also aligned the markers of linkage map LWA2010 to the Eggplant_V3 assembly and found that 832 (87.4%) markers could be assigned to the 12 pseudochromosomes (Table S4), which was less than that obtained using our data (99.4%). Generally, the pseudochromosomes showed a good collinearity between the new eggplant and Eggplant_V3 assemblies (Figure 1 and Table S5).

**Figure 1.**
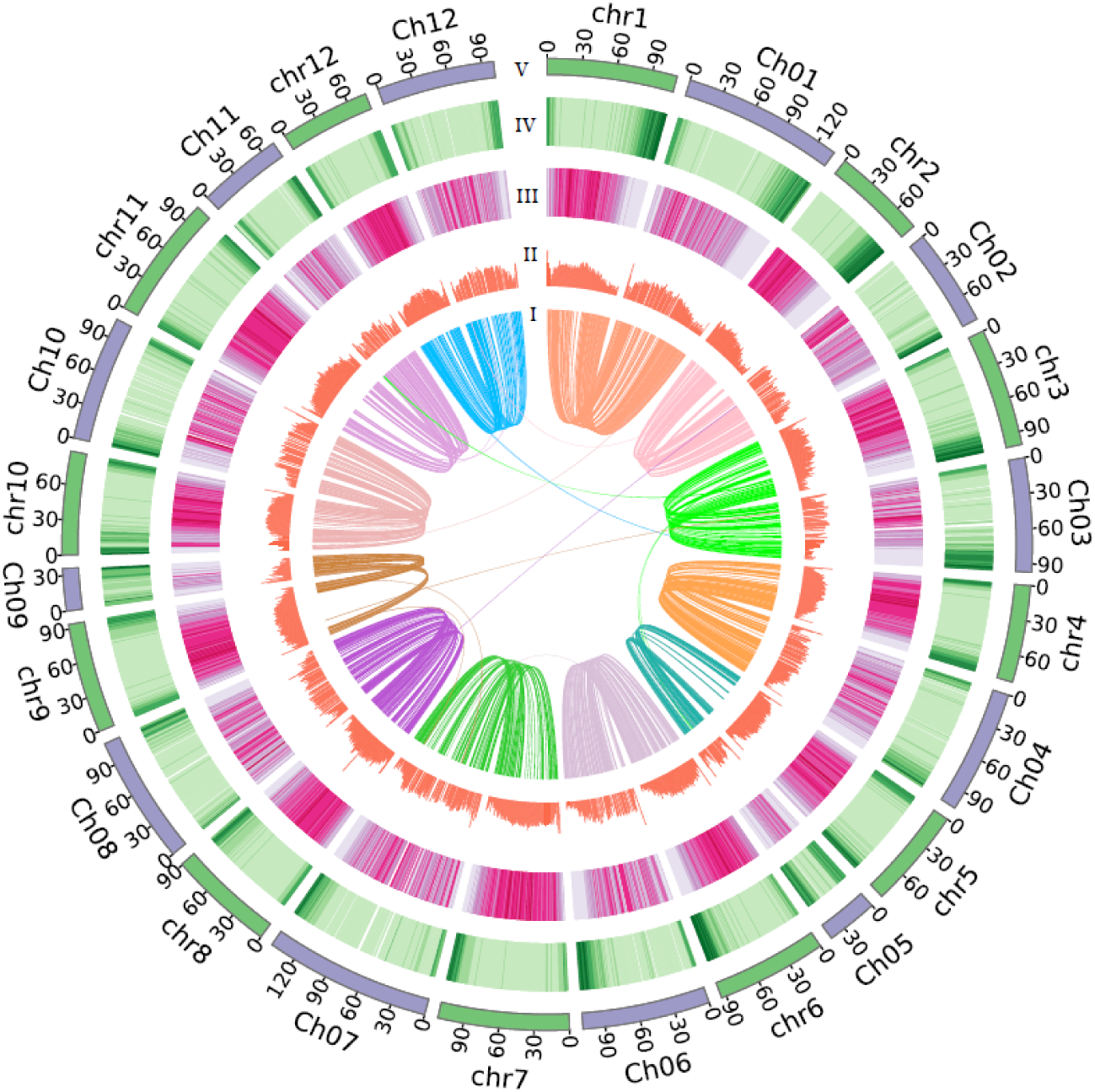
Comparison of the eggplant assemblies. I: Syntenic alignments between the new eggplant assembly and Eggplant_V3 assembly based on one-to-one orthologous genes processed by MCscan (Python version) with a C-score cutoff of 0.99 (links). II: GC content in non-overlapping 1-Mb windows (histograms). III: Percent coverage of transposable elements in non-overlapping 1-Mb windows (heat maps). IV: Gene density calculated on the basis of the number of genes in non-overlapping 1-Mb windows (heat maps). V: Lengths of pseudochromosomes (Mb) of the new eggplant assembly (green) and Eggplant_V3 assembly (purple).

A total of 70.1% of the assembly was annotated as repetitive sequences using a combination of homology-based and *de novo* approaches (Table S6). This proportion was consistent with that reported previously [28]. Transposable elements (TEs) play an important role in shaping eukaryotic genomes and driving their evolution [34]. In the eggplant, TEs accounted for 68.9% of the genome size, with long terminal repeats (LTRs) being the most predominant type (63.9% of the genome size) (Table S7). The proportions of TEs and LTRs were both less than those in the pepper [29, 35] and more than those in tomato [30] and potato [36]. The most abundant LTRs were the *Gypsy* elements (52%), followed by *Copia* (7.9%) (Table S7). This scenario was also observed in the sequenced pepper genome, indicating that the LTRs/*Gypsy* elements were the major driving force for the expansion of the eggplant genome. We then examined the insertion time of all LTRs based on sequence divergence. The eggplant appeared to have undergone a surge of retrotransposon amplification approximately 0.124 million years ago (Figure S3), suggesting that the expansion event was quite recent during its genome evolution.

To facilitate genome annotation of eggplant genes, we sequenced RNA samples from roots, stems, leaves, and flowers. The sequencing data were imported to the gene prediction pipeline, which also integrated homology-based and *de novo* strategies. We predicted 35,018 protein-coding genes, with an average gene length of 5,068 bp and an average of 4.7 exons per gene (Table S8). This number of genes is almost the same as that in tomato (35,768 genes), potato (39,028 genes), and pepper (35,845), indicating similar numbers of genes in this clade. The distribution of gene density was inversely correlated with TEs (Figure 1). BUSCO assessment of the predicted gene sets suggested 96.6% completeness, of which 94.2% and 2.4% were single-copy and duplicated genes, respectively (Table S9), suggesting the integrity of our new eggplant gene annotation. Further functional annotation using public databases indicated that 31,963 (91.3%) genes could be classified using at least one of the databases and 19,466 (55.6%) genes could be annotated using all five databases (Table S10). In addition, a total of 6,520 noncoding RNAs (ncRNAs) were found in the eggplant genome, including 116 microRNAs (miRNAs), 1,254 transfer RNAs (tRNAs), 4,629 ribosomal RNAs (rRNAs), and 521 small nuclear RNAs (snRNAs) (Table S11).

### Genome comparison and gene family evolution

Genome collinearity analysis of Solanaceae plants showed that some chromosomes were conserved; in particular, chromosomes 2, 6, and 7 retained a large percentage of collinear regions among eggplant, pepper, potato, and tomato (Figures 2a, S4).

**Figure 2.**
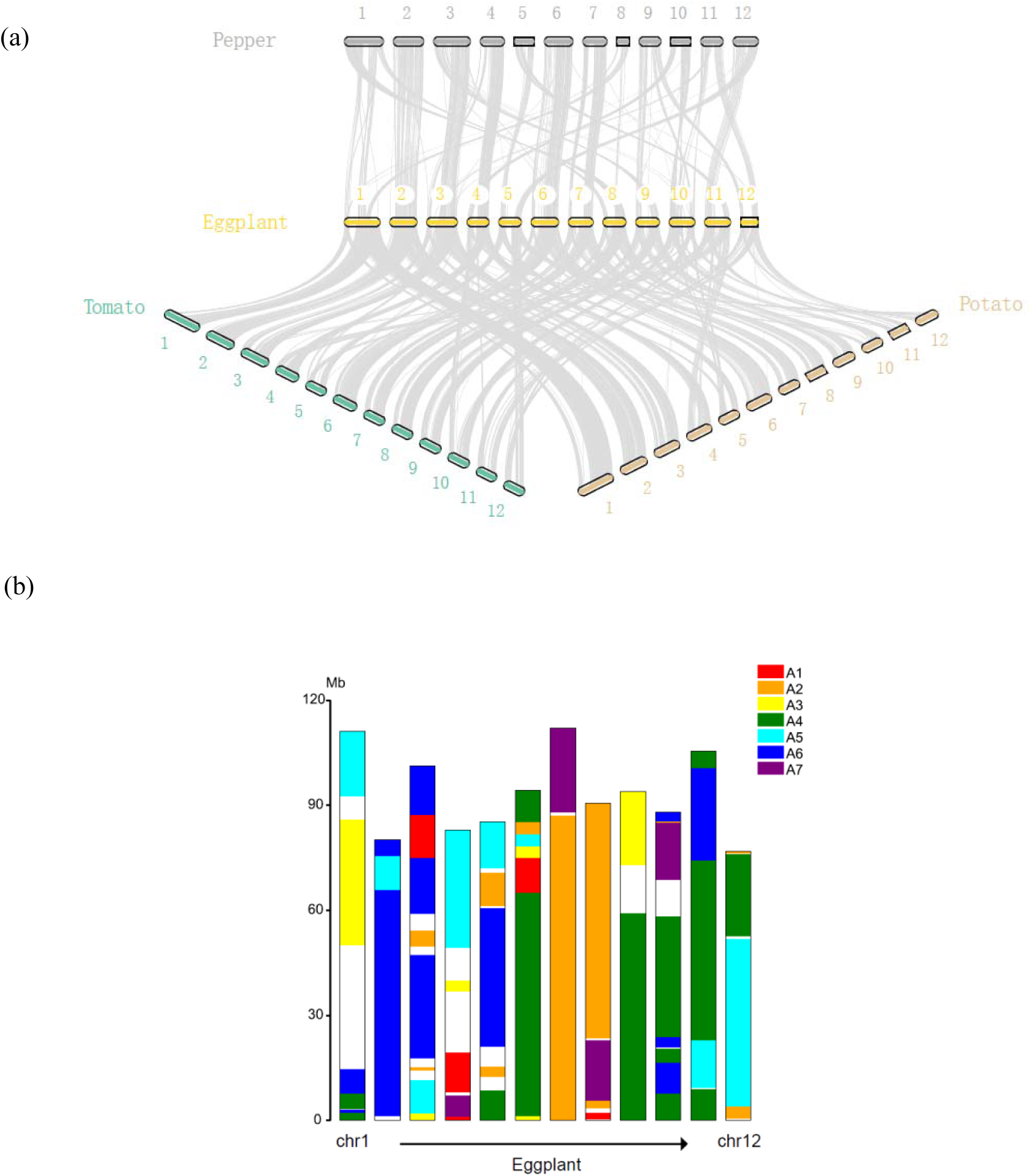
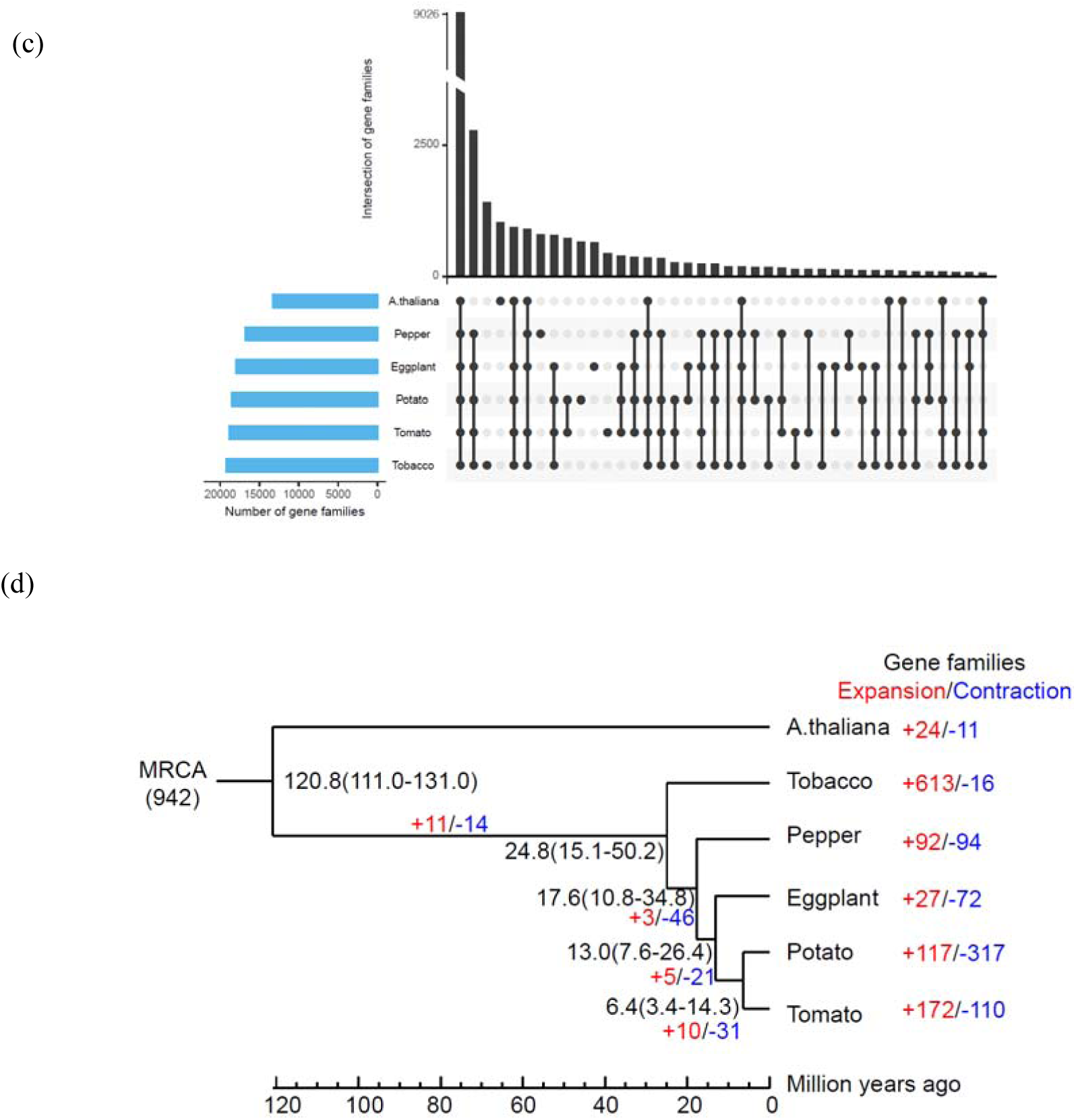
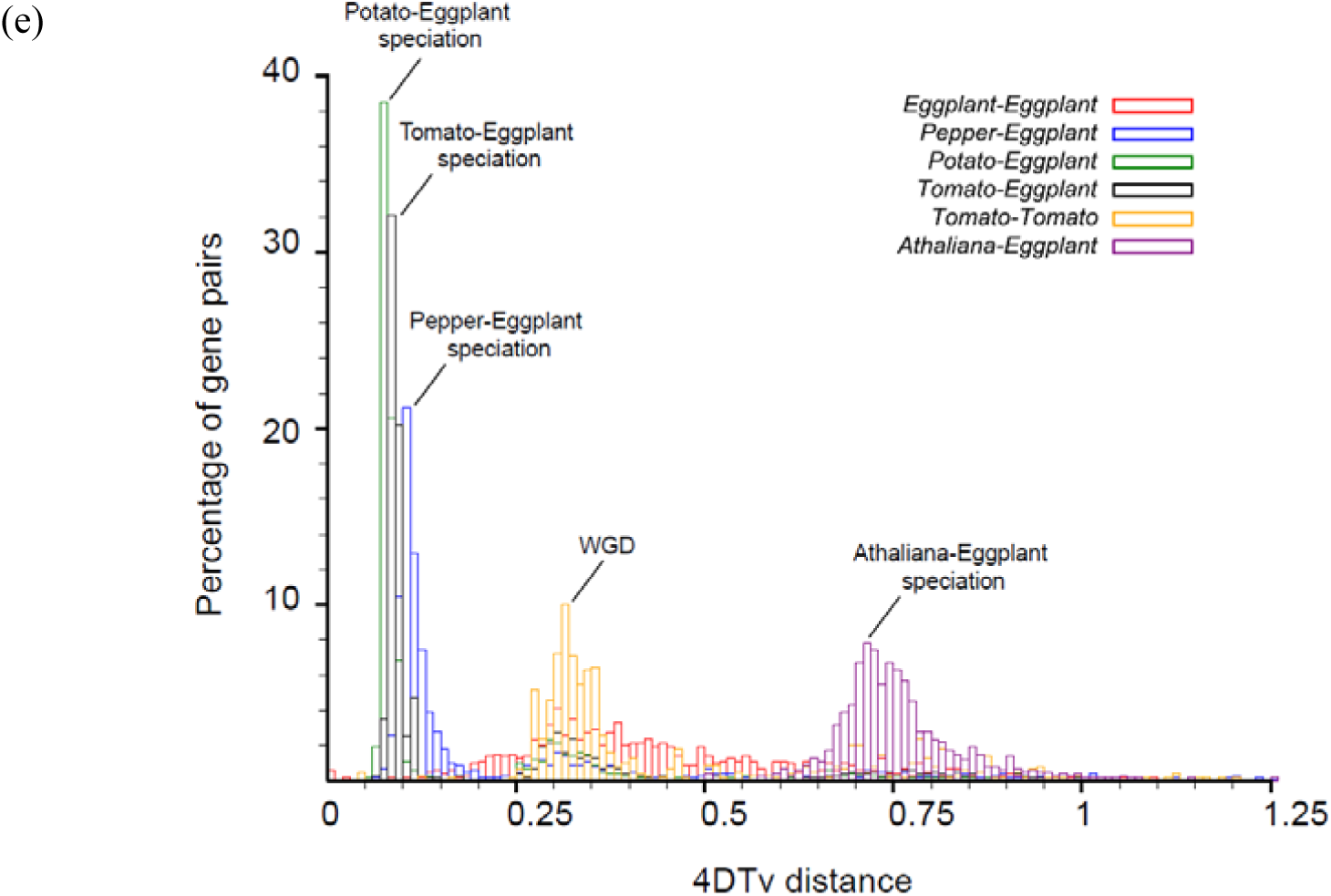
Comparative analysis and evolution of the eggplant genome. (a) Analysis of the synteny among Solanaceae genomes. Macrosynteny connecting blocks of >30 one-to-one gene pairs is shown. (b) Genome evolution of the eggplant from the ancestral eudicot karyotype (AEKPre-γ) of seven protochromosomes. Colors indicate the origin from the seven AEKPre-γ protochromosomes. White spaces represent chromosomal regions where ancestral origin was not assigned. (c) Intersections of gene families between six plant species (eggplant, pepper, potato, tobacco, tomato, and *Arabidopsis thaliana*). The figure was plotted using UpSetR [40], with the rows representing gene families and the columns representing their intersections. For each set that is part of a given intersection, a black filled circle is placed in the corresponding matrix cell. If a set is not part of the intersection, a light gray circle is shown. A vertical black line connects the topmost black circle with the bottommost black circle in each column to emphasize the column-based relationships. The size of the intersection is shown as a bar chart placed on top of the matrix so that each column lines up with exactly one bar. A second bar chart, showing the size of each set, is shown to the left of the matrix. (d) Phylogenetic tree with divergence times and history of orthologous gene families. Numbers on the nodes represent divergence times, with the error range shown in parentheses. The numbers of gene families that expanded (red) or contracted (blue) in each lineage after speciation are shown on the corresponding branch. MRCA, most recent common ancestor. (e) Genome duplication in Solanaceae genomes (pepper, tomato, potato, and eggplant) revealed by 4DTv analysis.

Based on the ancestral and lineage-specific whole-genome duplications reported for eudicots [37], we inferred genome evolution of the eggplant and other Solanaceae plants from the ancestral eudicot karyotype (AEKPre-γ) of seven protochromosomes. Figure 2b shows the chromosomes of the eggplant, with the seven protochromosomes of AEKPre-γ depicted in different colors. The map of the chromosomal regions that originated from different ancestral eudicot karyotypes (AEKs) is similar among eggplant, potato, and tomato (Figure S5 and Table S12) but much different from that of pepper. The pepper genome contains more predicted chromosomal regions, indicating that the genome of the pepper has undergone a much different process of genomic rearrangements to reach its current structure of 12 chromosomes, compared with that of the genomes of the other three solanaceous species.

We clustered the protein-coding genes of eggplant, pepper, potato, tobacco, tomato, and *Arabidopsis thaliana* into gene families (Table S13) and identified 25,620 gene families, of which 9,026 were shared by all six species. The intersections of the gene families are illustrated in Figure 2c. There are 358 gene families shared among the eggplant, pepper, potato, and tomato. In the eggplant, 26,596 genes were clustered into 17,926 gene families, of which 646 families were species-specific. Annotation of these specific genes showed various functions (Tables S14, S15), but they were particularly overrepresented in the chitin-related Gene Ontology (GO) categories. Chitin-binding genes are known as a pathogenesis-related gene family, which plays a fundamental role in the defense response of plants [38, 39]. This finding suggests possible response roles, related to biotic stress, in eggplant.

Analysis of evolution of gene families revealed that 27 gene families were expanded and 72 gene families were contracted in the eggplant (Figure 2d and Tables S16–S19). For the six plants, 799 single-copy genes were used to construct a phylogenetic tree and estimate their divergence times (Figure 2d). The data showed that the eggplant was separated from potato and tomato ∼12 million years ago during the Solanaceae evolution.

We then deduced whole-genome duplication (WGD) events in the eggplant based on the distribution of the distance-transversion rate at fourfold degenerate sites (4DTv methods) of paralogous gene pairs (Figure 2e). After the eggplant–*A. thaliana* speciation (peak at ∼0.71), there occurred a common Solanaceae WGD event (peak at ∼0.31). The divergence of eggplant–pepper occurred at a peak of ∼0.1, followed by eggplant–tomato (4dTv = 0.08) and eggplant–potato (4dTv = 0.07) divergence, which is consistent with the phylogenetic analysis. There is no evidence of an eggplant-specific WGD after the differentiation of *Solanum* plants.

In addition, we used the bidirectional best hit (BBH) method and recovered a total of 8,982 one-to-one orthologous gene sets among the five Solanaceae plants for positive selection gene (PSG) detection. In the eggplant, 364 PSGs were identified [*P* < 0.05, likelihood ratio test (LRT)], which were especially enriched in GO terms related to intermembrane lipid transfer (three PSGs), regulation of transcription, DNA-templated (24 PSGs), and DNA-binding transcription factor (TF) activity (16 PSGs) (Tables S20, S21).

### Identification of genes involved in disease resistance

In addition to a wide range of abiotic stresses such as the temperature, drought, and salt stress, eggplants are susceptible to a wide variety of biotic threats, including fungal pathogens and insect pests [41]. Most of the proteins encoded by the characterized resistance gene analogs (RGAs), including nucleotide-binding site (NBS)-containing proteins, receptor-like protein kinases (RLKs), and receptor-like proteins (RLPs), contain conserved domains, such as NBS, leucine-rich repeat (LRR), and Toll/interleukin-1 receptor (TIR) [42]. Using a genome-wide scanning pipeline [43], we identified 1,023 RGAs in the eggplant (Table S22), which was comparable to the number of RGAs in tomato, slightly lower than that in potato, and much lower than that in the pepper (Table 2). Pepper contains almost twice the total number of RGAs in each of the three *Solanum* spp., consequent to tandem duplication of genes, which also resulted in its genome expansion [29]. Half of RGAs in the eggplant belonged to the RLK category, and there were 285 NBS-related RGAs, of which 33 were of the TIR type. We noticed that over 80% of RGAs clustered near the head and tail of chromosomes, and this distribution pattern was consistent with the overall gene distribution in the eggplant genome.

**Table 2.** Comparison of RGAs among four Solanaceae genomes. NBS, nucleotide-binding site; CC, coiled-coil; LRR, leucine-rich repeat; TIR, Toll/interleukin-1 receptor; TM, transmembrane; RLK, receptor-like kinase; RLP, receptor-like protein; CNL, CC-NBS-LRR; TNL, TIR-NBS-LRR; CN, CC-NBS; TN, TIR-NBS; NL, NBS-LRR; TX, TIR-unknown domain; Others, CC-TIR.

| Species | NBS encoding |  |  |  |  |  |  |  | RLP | RLK | TM-CC | Total |
| --- | --- | --- | --- | --- | --- | --- | --- | --- | --- | --- | --- | --- |
|  | NBS | CNL | TNL | CN | TN | NL | TX | Others |  |  |  |  |
| Eggplant | 82 | 65 | 21 | 18 | 12 | 75 | 11 | 1 | 84 | 511 | 143 | 1,023 |
| Tomato | 64 | 66 | 22 | 13 | 9 | 83 | 13 | 1 | 87 | 533 | 148 | 1,039 |
| Potato | 100 | 90 | 35 | 33 | 12 | 148 | 30 | 4 | 156 | 562 | 111 | 1,281 |
| Pepper | 282 | 137 | 19 | 75 | 15 | 238 | 19 | 7 | 203 | 687 | 151 | 1,833 |

There were 15 RGAs overlapped with PSGs, including nine RLK-encoding RGAs, three encoding transmembrane coiled-coil-containing proteins, two encoding NBS-LRR-containing proteins, and one encoding a TIR-NBS-LRR-containing protein (Table S23). Among these, eight genes could be assigned to known resistance genes using the reference PR proteins from the latest PRGdb [44]. We inferred that these positively selected resistance genes probably played a fundamental role in eggplant self-defense. Further mining revealed an interesting orthoMCL group (129 genes), whose analysis indicated explosive gene expansion in eggplant (21 genes) and pepper (96 genes), in contrast to tomato (three genes) and potato (two genes). Tobacco had seven members in this group, while *Arabidopsis* did not have any. All of these genes were annotated using PRGdb as encoding bacterial spot resistance gene *BS2* (Table S24) [45]. In a maximum-likelihood phylogenetic tree, constructed using IQ-TREE [46], the 21 eggplant genes formed a monophyletic cluster (Figure S3) and, moreover, were found to be tandemly clustered at the head of chromosome 12. We inferred that the occurrence of these genes might be a consequence of tandem duplication events during eggplant genome evolution, which was also observed in pepper [29].

### Identification of genes involved in CGA synthesis

CGAs (esters of certain *trans*-cinnamic acids and quinic acid) are major phenolic metabolites in the eggplant, which typically account for 80% to 95% of total hydroxycinnamic acids in the fruit flesh [47, 48]. CGAs play a role in plant defense and as antioxidants and are accumulated in many Solanaceae plants [47, 49]. However, the CGA content in the eggplant has been reported to be roughly 10 and 100 times higher than that in tomato and potato, respectively [50]. CGA is well known to be beneficial for human health, mainly owing to its antioxidant, anti-inflammatory, antipyretic, anticarcinogenic, antimicrobial, analgesic, neuroprotective, cardioprotective, hypotensive, anti-obesity, and antidiabetic properties [48, 51]. Moreover, CGA is highly stable at high temperatures, and its content increases after cooking [52]. Thus, eggplant is considered to be the best source of CGA among the Solanaceae.

The biosynthesis of CGA occurs in eggplants through the phenylpropanoid pathway, which involves six key enzymes [47, 53]. The three initial steps, catalyzed by phenylalanine ammonia-lyase (*PAL*), cinnamate 4-hydroxylase (*C4H*), and 4-coumaroyl-CoA ligase (*4CL*), produce the intermediate *p*-coumaroyl-CoA (Figure 3a). Using homologous gene comparison, we identified six *PAL*, two *C4H*, and five *4CL* candidate genes in the eggplant genome (Figures 3b, S7 and Table S25). *Arabidopsis* contains four *PAL* genes, two of which (*AtPAL1* and *AtPAL2*) are associated with lignin and flavonoid biosynthesis [54]. Three eggplant *PAL* genes were in three distinct phylogenetic groups, and the other three clustered together, while the four *Arabidopsis PAL* genes formed a single clade (Figure S8). Overexpression of *AtPAL2* in tobacco resulted in a twofold increase in the CGA content [55]. C4H is a cytochrome P450 (CYP) monooxygenase from the CYP73A subfamily, and only one member, designated CYP73A5, exists in *Arabidopsis*. One *C4H* gene (EGP13151) in eggplant exhibited more sequence identity with the *Arabidopsis* gene than did the other (EGP24021) (86% versus 65%, respectively). Missense mutations in *C4H* result in metabolic changes, threatening plant survival [54, 56]. Downregulation of *C4H* resulted in a decrease of CGA levels in tobacco, as well as in a feedback inhibition of *PAL* activity [57]. It has been reported that *Arabidopsis* contains four *4CL* genes, two of which are involved in lignin biosynthesis, one is related to flavonoid biosynthesis, and the last one preferentially towards erulate and sinapate instead of 4-coumarate [54]. The eggplant has five *4CL* genes, which is similar to the number in the other three Solanaceae members but is only half of that in tobacco (Figure 3b). Phylogenetic analysis revealed that each *4CL* was in a distinct clade (Figure S9). A previous study has shown that the expression levels of *PAL, C4H*, and *4CL* in eggplants at the commercially ripe stage were notably higher in the fruit flesh and skin than in other tissues, indicating their correlation with the higher CGA content in the fruit [50].

**Figure 3.**
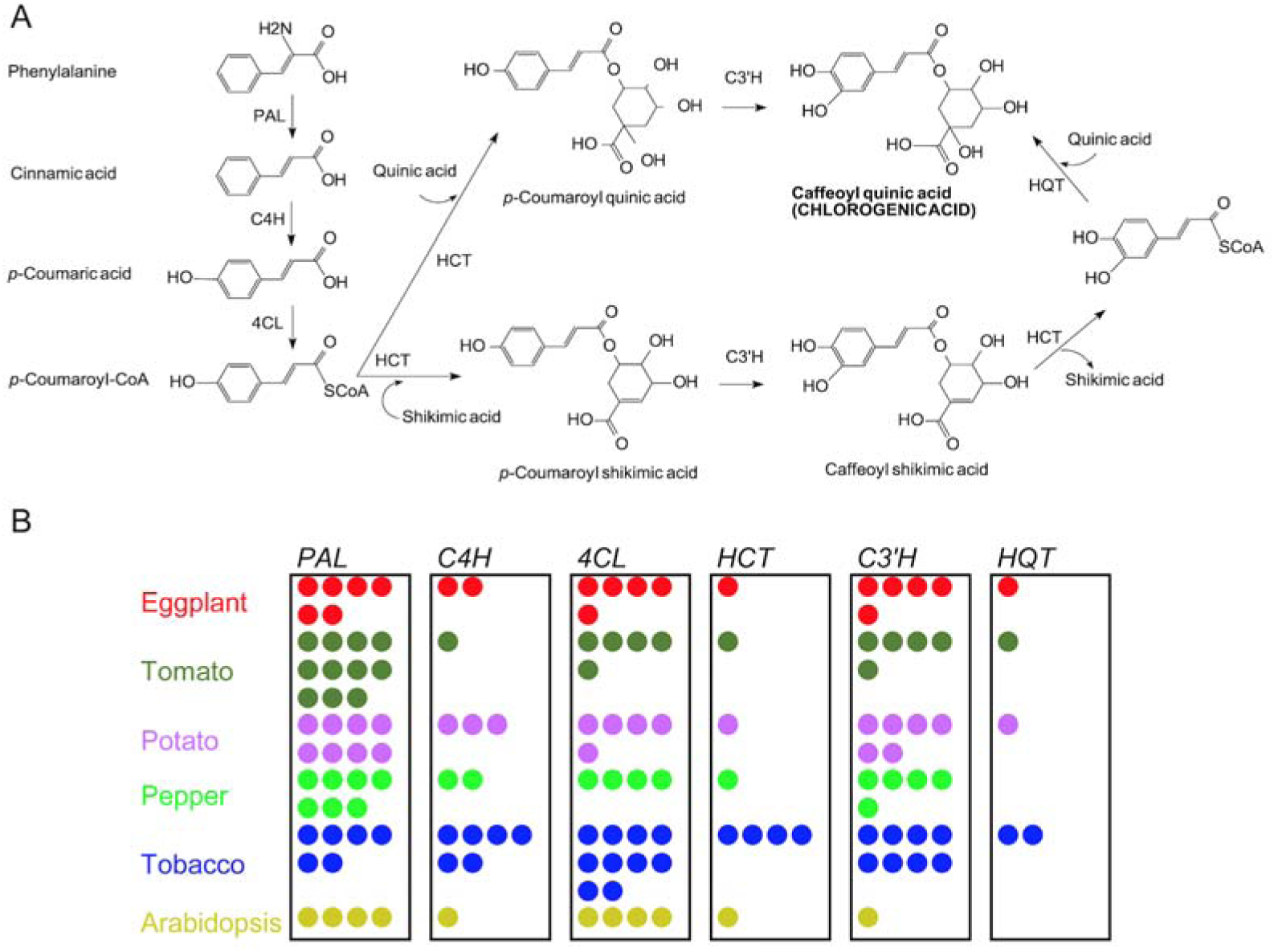
Genes involved in chlorogenic acid (CGA) synthesis. (a) Biochemical pathway for CGA synthesis in the eggplant. The enzymes involved are as follows: *PAL*, phenylalanine ammonia-lyase; *C4H*, cinnamate 4-hydroxylase; *4CL*, 4-coumaroyl-CoA ligase; *HCT*, hydroxycinnamoyl-CoA:shikimate hydroxycinnamoyl transferase; *C3*′*H, p*-coumaroyl ester 3′-hydroxylase; *HQT*, hydroxycinnamoyl-CoA:quinate hydroxycinnamoyl transferase. (b) Orthologous genes involved in CGA biosynthesis from eggplant (red), tomato (green), potato (purple), pepper (light green), tobacco (blue), and *Arabidopsis* (yellow), identified using orthoMCL, followed by manual inspection. Each circle represents one gene.

After the three initial steps in CGA biosynthesis, two possible pathways have been suggested (Figure 3a): (1) *p*-coumaroyl-CoA is converted into *p*-coumaroyl quinic acid with quinic acid via hydroxycinnamoyl-CoA:shikimate hydroxycinnamoyl transferase (*HCT*), followed by hydroxylation to form CGA via *p*-coumaroyl ester 3′-hydroxylase (*C3*′*H*); and (2) *p*-coumaroyl-CoA is converted into *p*-coumaroyl shikimic acid with shikimic acid via *HCT*, followed by hydroxylation to form caffeoyl shikimic acid via *C3*′*H*. Caffeoyl shikimic acid, catalyzed by *HCT*, is converted into caffeoyl-CoA, which is then converted into CGA by *trans*-esterification with quinic acid via hydroxycinnamoyl-CoA:quinate hydroxycinnamoyl transferase (*HQT*) [58]. *HCT* and *HQT* are closely related BAHD-like acyltransferases [59, 60], and both are encoded by single-copy genes in eggplant, tomato, and potato (Figure 3b and Table S25). However, *HQT* is absent in *Arabidopsis* and pepper. Overexpression of *HQT* in *AtPAL2*-overexpressing tobacco plants resulted in a 1.4-fold increase in the CGA content, while silencing of *HQT* resulted in a ∼50% reduction in CGA [55]. In tomato, overexpression of *HQT* led to an increase in CGA accumulation, improving the plant antioxidant capacity and bacterial pathogen resistance [61]. RNAi suppression of *HQT* in potato resulted in a ∼90% reduction in CGA and early flowering [62]. In the eggplant, the expression of *HQT* was the strongest in the fruit flesh and skin, compared with that in other tissues at the ripe stage [50]. *C3*′*H* is a CYP monooxygenase belonging to the CYP98A subfamily; in *Arabidopsis* [63], *C3*′*H* (designated CYP98A3) is one of three members of this family (the other two members are AT1G74540-CYP98A8 and AT1G74550-CYP98A9). Unlike *Arabidopsis*, multiple homologs of *C3*′*H* were detected in the five Solanaceae species, including five *C3*′*H* genes in the eggplant. Similar to *4CL*, each *C3*′*H* was located in a distinct phylogenetic clade (Figure S10). We inferred that these gene duplications had evolved via independent processes, which led to divergent gene functions or neofunctionalization, responsible for the remarkable increase of CGA biosynthesis in the eggplant.

Polyphenol oxidases (PPOs), which oxidize specific phenolic substrates released from vacuoles upon tissue damage to highly reactive quinones, play key roles in plant defense mechanisms against pests and pathogens [64, 65]. However, oxidation of these high-level phenolics, including CGA, results in flesh browning, which negatively affects the apparent quality of eggplants [48]. In this respect, simultaneous breeding for a high CGA content and low PPO activity would result in cultivars with better fruit quality and reduced flesh browning [47]. We identified nine PPOs in the eggplant, with eight genes tandemly clustered at the end of chromosome 8 and one located on chromosome 2 (Table S26 and Figure S7). Previously published studies discovered six PPO genes in the eggplant [65], and five, except *PPO6*, could be anchored to chromosome 8 using a linkage map [48]. Protein sequence identities ranged from 92% to 99% when comparing these six genes to our dataset (Table S27). We further examined PPOs in other species. There were nine, eight, eight, and twelve PPO homologs in tomato, potato, pepper, and tobacco, respectively (Figure S11). The absence of *PPO*s in *Arabidopsis* has been discussed [66]. We also observed that the distribution patterns of PPO genes in the tomato and potato genomes were highly similar to that in the eggplant genome, with one located on chromosome 2 and the rest clustered at the end of chromosome 8 (Table S26), indicating a highly conserved synteny among the three solanaceous species.

### Identification of genes encoding transcription factors

Plant secondary metabolism is regulated by TFs, which act as transcriptional activators or repressors [67, 68]. We identified 1,702 TF-encoding genes in the eggplant, representing 4.86% of the total genes. The number of members from each TF family in the eggplant was comparable to that in four other plants but was much lower than that of certain families in tobacco, such as bHLH, ERF, and NAC (Table S28). Genes encoding MYB TFs, containing conserved MYB DNA-binding domains, are a large family of functionally diverse genes, which can be classified into four subfamilies, 1R, R2R3, 3R, and 4R [67]. The R2R3 subfamily is the largest and considered to comprise the major phenylalanine-derived compound modulators in plants. We identified 121 MYB and 61 MYB-related TFs in the eggplant, of which 112 belonged to the R2R3 subfamily, and most of them could be categorized into 20 subgroups (Table S29) according to the previously characterized R2R3 genes in *Arabidopsis* [69, 70]. Several subgroups (SG4–SG7) have been found to regulate the phenylpropanoid pathway, including anthocyanin and flavonol biosynthesis [67]. We identified three SG3, four SG4, three SG6, and three SG7 genes in the eggplant. The *SmMyb1* gene, belonging to SG6, was reported to regulate CGA accumulation and anthocyanin biosynthesis [50]. No SG5 members were identified based on the current criteria. We also found a gene cluster, which was located at the end of chromosome 7 and contained five members, four belonging to SG2 and one belonging to SG3, suggesting their key roles in regulating self-defense [71, 72].

## Conclusion

We sequenced and assembled the genome of the eggplant and greatly improved the quality and integrity of the sequence compared with those of previously published draft sequences. As a vital crop in the Solanaceae, eggplants are cultivated and consumed worldwide. However, there have been much fewer studies of the eggplant than of other members of the Solanaceae, such as tomato and potato, which have been established as biological models for studying the development of fleshy fruits and tubers, respectively. The main reason is due to the lack of a high-quality reference genome of the eggplant. Although a genome sequence of the inbred eggplant line ‘67/3’ has been published recently, our assembly showed several advantages, including a longer contig N50 (5.3 Mb vs. 16.7 kb), fewer total scaffolds (319 vs. 10,383), and a much smaller size of gaps (0.003% vs. 28.23%). Genome validation using a linkage map confirmed a high accuracy of our assembly.

We comprehensively characterized genes involved in disease resistance, CGA synthesis, and polyphenol oxidation, as well as those encoding TFs, thus demonstrating a significant value of the reference genome sequence. We also conducted comparative analysis of the eggplant genome with those of four other species of the Solanaceae and *Arabidopsis*. This study will facilitate the breeding of eggplant cultivars with strong disease resistance, high nutritional value, and low browning.

## Methods

### Sample preparation

Guiqie1 (*S. melongena*) plants were collected from the Vegetable Research Institute, Guangxi Academy of Agricultural Science (28°N and 118°E), Guangxi province, China. Roots, stems, leaves, and flowers of Guiqie1 were harvested, immediately frozen in liquid nitrogen, and stored at −80 °C until use. Genomic DNA was isolated from leaf tissues using the DNeasy plant mini kit (Qiagen). RNA was extracted using the RNeasy plant mini kit (Qiagen).

### DNA sequencing

#### Illumina short-read sequencing

Purified DNA was sheared using a focused ultrasonicator (Covaris) and then used for 350-bp paired-end library construction with the Next Ultra DNA library prep kit (NEB) for Illumina sequencing. Sequencing was performed on the Illumina NovaSeq platform.

#### SMRT long-read sequencing

SMRTbell DNA libraries (∼20 kb) were prepared using the BluePippin size selection system following the officially released PacBio protocol. Long reads were generated using the PacBio Sequel system.

#### Hi-C library construction and sequencing

A Hi-C library was prepared using the Dovetail Hi-C library preparation kit. Briefly, nuclear chromatin was fixed in young eggplant seedlings with formaldehyde and extracted. Fixed chromatin was digested with *Dpn*II, and sticky ends were filled in with biotinylated nucleotides and ligated. Then, crosslinks were reversed, and purified DNA was treated to remove any free biotin from ligated fragments. DNA was then sheared to a size of ∼350 bp, and biotinylated fragments were enriched through streptavidin bead pulldown, followed by PCR amplification to generate the library. The library was sequenced on the Illumina NovaSeq platform.

### Genome assembly and evaluation

A diploid contig assembly of the eggplant genome was carried out using FALCON, followed by FALCON-Unzip, integrated in the pb-assembly tool suite (v0.0.4). The resulting assembly contained primary contigs (partially phased haploid representation of the genome) and haplotigs (phased alternative alleles for a subset of the genome). Two rounds of contig polishing were performed. For the first round, as part of the FALCON-Unzip pipeline, primary contigs and secondary haplotigs were polished using haplotype-phased reads and the Quiver consensus caller. For the second round of polishing, we concatenated the primary contigs and haplotigs into a single reference and then mapped all raw reads to the combined assembly reference using pbmm2 (v0.12.0), followed by consensus calling with Arrow (GenomicConsensus v2.3.3). After a draft set of contigs was generated, the Dovetail Hi-C kit was run for Hi-C-based scaffolding with cloud-based HiRise software [73]. Finally, Pilon (v1.22) was used to correct errors introduced into the assembly from long reads.

To assess the completeness of the assembled eggplant genome, we performed BUSCO analysis by searching against the conserved 1,440 Embryophyta gene set (v3.0, lineage dataset embryophyta_odb9).

### Repeat annotation

Tandem repetitive sequences were identified within the eggplant genome using Tandem Repeats Finder (v4.07). The interspersed repeats were determined using a combination of homology-based and *de novo* approaches. The homology-based approach, with the RepBase (v21), was used to identify TEs by searching against the eggplant genome assembly at the DNA and protein levels using RepeatMasker (v4.0.7; http://www.repeatmasker.org/) and ProteinRepeatMask (v4.0.7), respectively. A *de novo* repeat library was customized using RepeatModeler (v1.0.8) and LTR_FINDER (v1.0.6) [74] and then imported to RepeatMasker to identify repetitive elements. Additionally, the results from LTR_FINDER were integrated, and false positives were removed from the initial predictions using the LTR_retriever pipeline [75]. The insertion time was estimated as T = K/2μ, where K is the divergence rate, and μ is the neutral mutation rate. A neutral substitution rate of 9.6 × 10^−9^ was used for the eggplant [76].

### Gene annotation

Protein-coding gene predictions were conducted through a combination of homology-based, *de novo*, and transcriptome-based prediction methods. Proteins for six plant genomes (*A. thaliana, C. annuum, S. tuberosum, N. tabacum, S. lycopersicum, and S. melongena* SME_r2.5.1) were downloaded from Phytozome (release 13), the National Center for Biotechnology Information (NCBI), and the Eggplant Genome DataBase. Protein sequences were aligned to the assembly using genblasta (v1.0.4). GeneWise (v2.4.1) was used to predict the exact gene structure of the corresponding genomic regions on each genblasta hit. Three *ab initio* gene prediction programs, Augustus (v3.2.1), GlimmerHMM (v3.0.4), and SNAP (v2006-07-28), were used to predict coding regions in the repeat-masked genome. Finally, RNA-seq data were mapped to the assembly using hisat2 (v2.0.1); stringtie (v1.2.2) and TransDecoder (v3.0.1) were then used to assemble the transcripts and identify candidate coding regions in gene models. All gene models predicted by the above three approaches were combined using EvidenceModeler into a non-redundant set of gene structures. The produced gene models were finally refined using PASA v2.3.3. Functional annotation of protein-coding genes was achieved using BLASTP (E-value: 1e−05) against two integrated protein sequence databases, SwissProt and TrEMBL. Protein domains were annotated using InterProScan (v5.30). The GO terms for each gene were extracted with InterProScan. The pathways in which genes might be involved were assigned using BLAST against the KEGG database (release 84.0), with an E-value cutoff of 1e−05.

Four types of ncRNAs, namely, miRNAs, tRNAs, rRNAs, and snRNAs, were annotated. The tRNA genes were predicted using tRNAscan-SE (v1.3.1). The rRNA fragments were predicted through alignment to *Arabidopsis* and rice template rRNA sequences using BlastN (v2.2.24), with an E-value of 1e−5. The miRNA and snRNA genes were determined by searching against the Rfam database (release 12.0) using INFERNAL (v1.1.1).

### Genome comparison and gene family and phylogenetic analyses

The AEK genes in the modern genome of the grape were obtained from Murat et al. [37]. Based on genome alignments using the cumulative identity percentage and cumulative alignment length percentage BLAST parameters [77], we identified homologous genes of AEK in the modern genomes of Solanaceae plants. Synteny blocks between the genomes of Solanaceae plants were detected using the GRIMM-Synteny software (http://grimm.ucsd.edu/GRIMM/), with groups of fewer than five genes filtered out; then, the synteny blocks were assigned to the seven protochromosomes based on the homologous genes of AEK.

OrthoMCL (v2.0.9) [78] was used to cluster gene families from *A. thaliana, C. annuum, S. tuberosum, N. tabacum, S. lycopersicum*, and *S. melongena*. CAFÉ (v3.1) [79] was used to determine gene family expansion and contraction.

A total of 799 single-copy genes were used to construct a phylogenetic tree for the six plant genomes. Fourfold degenerate sites were extracted from each family and concatenated to form one supergene for each species. The GTR-gamma substitution model was selected, and PhyML (v3.0) [80] was used to reconstruct the phylogenetic tree. The divergence times among the six plants were estimated using the MCMCtree program (v4.4) as implemented in the Phylogenetic Analysis of Maximum Likelihood (PAML) package, with an independent rate clock and the JC69 nucleotide substitution model. The calibration times of divergence between *A. thaliana* and *S. lycopersicum* (111–131 million years ago) were obtained from the Time Tree database [81].

To detect PSGs in the eggplant genome, one-to-one orthologs were identified among the six plants using BLASTP, based on the BBH method with a sequence coverage >30% and identity >30%, followed by selection of the best match. A total of 8,982 one-to-one orthologous gene sets were found among *C. annuum, S. tuberosum, N. tabacum, S. lycopersicum*, and *S. melongena*. The branch-site model incorporated in the PAML package was used, with the eggplant used as the foreground branch and pepper, potato, and tomato used as background branches. The null model used in the branch-site test assumed that the Ka/Ks values for all codons in all branches were ≤1, whereas the alternative model assumed that the foreground branch included codons evolving at Ka/Ks >1. A maximum LRT was used to compare the two models. The *P*-value was calculated using the chi-squared distribution with one degree of freedom, and then *P*-values were adjusted for multiple testing using the false discovery rate (FDR) method. Genes were identified as positively selected when FDR was <0.05. Furthermore, we required that at least one amino acid site possessed a high probability of being positive selected (Bayes probability >95%). If no amino acid in PSG passed this cutoff, such gene was identified as false positive and excluded. GO enrichment was derived using Fisher’s exact test and adjusted using the Benjamini–Hochberg method with the cutoff set at *P* < 0.05.

### Identification of disease resistance genes

The RGAugury pipeline (https://bitbucket.org/yaanlpc/rgaugury) [43] was used to screen the entire gene set for RGA prediction. The default *P*-value cutoff for initial RGA filtering was set to le−5 for BLASTP.

### Identification of CGA synthesis-related genes and phylogenetic analysis

To identify CGA synthesis-related genes, homologous *Arabidopsis* genes were mined from the literature and downloaded. Corresponding gene family results were extracted and manually inspected. HMMER or BLASTP were used whenever necessary. Protein sequences were aligned using muscle (v3.8.31). Maximum-likelihood phylogenetic trees were constructed using IQ-TREE (v1.6.11), with 1,000 bootstrap replicates, and further illustrated in MEGA (v7.0.26).

### Identification and classification of TFs

The Plant Transcription Factor Database v5.0 (planttfdb.cbi.pku.edu.cn) was used to identify TFs [82]. R2R3-MYB TFs were further characterized using the corresponding members in *Arabidopsis* [67, 69], and motifs were verified using MEME (v5.0.5) [83]. Subgroups were designated as previously reported [67, 70].

## Availability

The genome assembly and the sequencing data used for *de novo* whole-genome assembly are available from the China National GeneBank (CNGB) Nucleotide Sequence Archive (CNSA) under accession number CNP0000734.

**Supporting information**

Additional supporting information may be found online in the Supporting Information section at the end of the article.

